# Metabolic Reprogramming Driven by *mpSte11A* Deletion Redirects Carbon Flux toward Overproduction of *Monascus* Pigments in *Monascus* spp

**DOI:** 10.64898/2026.02.02.703344

**Authors:** Tingya Wang, Yali Duan, Yuting Liu, Mu Li

**Affiliations:** College of Food Science and Technology, Huazhong Agricultural University, Hubei Province, Wuhan 430070, China; Key Laboratory of Environment Correlative Dietology, College of Food Science and Technology, Hubei International Scientific and Technological Cooperation Base of Traditional Fermented FoodsHuazhong Agricultural UniversityHubei Province, Wuhan 430070, China

**Keywords:** Carbon flux, Serine/threonine protein kinases, STE11, *Monascus* spp., Fungal cell factory

## Abstract

*Monascus* spp., a food fermentation microorganism, produced valuable secondary metabolites including *Monascus* pigments (MPs) which served as natural food colorants. However, rational metabolic engineering to enhance MPs production remained limited by the lack of regulatory targets that govern metabolic branching. Mitogen-activated protein kinase cascades, particularly the STE20-STE11-STE7 core module, regulated fungal growth and metabolism, but their roles in MPs biosynthesis remain unexplored. In this study, we functionally characterized MpSte11A, the first STE11 homolog identified in *Monascus* spp., through bioinformatic analysis and genetic manipulation. Most importantly, deletion of *mpSte11A* triggered a profound metabolic shift, which resulted in a 22-fold increase in MPs production. Integrated transcriptomic and metabolomic analysis revealed that MpSte11A functioned as a metabolic gatekeeper where its deletion redirected carbon flux from primary to MPs biosynthesis by controlling the TCA cycle. These findings not only elucidated the signaling role of the MAPK cascade in *Monascus* spp. specialized metabolism but also provided a robust strategy for re-engineering carbon partitioning to maximize the output of high-value secondary metabolites in filamentous fungal cell factories.

**Importance:** Filamentous fungi are versatile cell factories for the production of diverse high-value secondary metabolites, but the rational enhancement of these compounds is often limited by a lack of universal regulatory targets. In this study, we employed the food-fermentation fungus *Monascus* spp. as a model and identified MpSte11A as a master “metabolic gatekeeper” that governed the trade-off between fungal growth and secondary metabolism. By disrupting this single signaling node, we achieved a remarkable 22-fold increase in compound production. This significant enhancement resulted from a systematic redirection of carbon flux from primary growth (the TCA cycle) to secondary biosynthesis. This work provided a precise molecular blueprint for the reprogramming of fungal metabolism. It also demonstrated that the tuning of core MAPK modules is a powerful and broadly applicable strategy for the engineering of robust fungal cell factories producing a wide array of bioproducts.

**Graphical Abstract (For Table of Contents Only):** 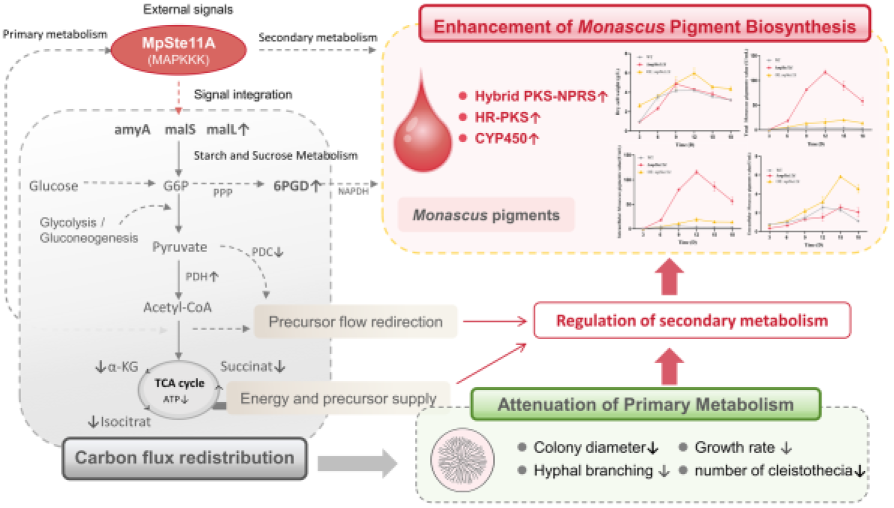

## Introduction

Filamentous fungi serve as major sources of industrial compounds and represent the cornerstone for producing secondary metabolites and many enzymes(1, 2). *Monascus* spp., a food fermentation microorganism, produces *Monascus* pigments (MPs) as natural food colorants(3, 4). Globally, approximately one billion people consume foods containing MPs in their daily diets(5). Considerable progress has been made in enhancing MPs production through strategies such as overexpressing biosynthetic genes or eliminating competing pathways(6, 7). However, these approaches often yielded modest improvementsa because they failed to address an issue where regulators controlling global metabolic flux distribution between primary growth and secondary metabolism are unclear.

Mitogen-activated protein kinase (MAPK) cascades are distributed in fungi and integrate external signals to regulate growth and metabolism(8, 9). Serine/threonine protein kinases, which are key components of multiple signaling pathways, function as evolutionarily conserved centers in complex signaling networks(10–12). Within the MAPK cascade, the STE20-STE11-STE7 module serves as the central regulatory axis. It integrates signals from diverse upstream pathways, including the high-osmolarity glycerol, invasive growth, cell wall integrity and pheromone response pathways, to regulate cellular growth, differentiation, and metabolism(13–15). This module not only regulates developmental programs but also controls resource allocation between primary and secondary metabolism, making it an attractive target for metabolic engineering.

Within the STE20-STE11-STE7 module, STE11 holds a uniquely central role as a shared MAPKKK, integrating signals from multiple upstream kinases (such as STE20) and relaying them to pathway-specific MAPKKs(16–22). This made STE11 an ideal target for regulating multiple MAPK pathways. Although STE11 has been widely studied in pathogenic fungi and model yeasts(23, 24), its roles in filamentous fungi remained poorly understood. This knowledge gap is especially critical for *Monascus* spp. because no STE11 homolog has been functionally characterized despite its potential as a major regulator linking growth and MPs biosynthesis.

In this study, we functionally characterized *mpSte11A*, the first STE11 homolog in *Monascus* spp., through genetic manipulation, phenotypic profiling, and integrated multi-omics analysis. Notably, deletion of *mpSte11A* dramatically enhanced MPs production by 22-fold compared to the wild-type strain. Integrated transcriptomic and metabolomic profiling revealed that MpSte11A functioned as a metabolic gatekeeper that control carbon flux distribution. Its absence redirected carbon flux from growth toward MPs biosynthesis through TCA cycle suppression and biosynthetic pathway rebalancing. These findings demonstrated that MpSte11A served as a key regulator linking developmental programs with MPs biosynthesis and also offered a target for metabolic engineering in other fungi.

## Results

### Bioinformatic Analysis Revealing a Candidate STE11 Homolog in *Monascus* spp

To date, no STE11 homolog has been functionally characterized in *Monascus* spp. Using STE11 protein sequences previously reported in fungi as queries(25), BLAST analysis identified 76 putative STE11-like proteins in *Monascus* ruber M7. Among these proteins, MpSte11A exhibited the highest coverage of 93%, with an identity of 36.7% and an E-value of 1×10^-22^..

We constructed a phylogenetic tree based on MpSte11A and the STE11 proteins derived from different fungal subphyla to analyze their phylogenetic relationships. (Fig. 1A). The analysis showed that MpSte11A is most closely related to the STE11 proteins of *Monascus purpureus.* Phylogenetic reconstruction resolved five major groups: *Zygomycotina, Ascomycotina, Basidiomycotina, Mastigomycotina*, and *Saccharomycotina*. MpSte11A clustered within the Ascomycotina clade, suggesting potential functional similarity to STE11 proteins from other ascomycetous fungi. Collectively, these results indicated that MpSte11A was highly conserved across fungi.

**FIG 1.**
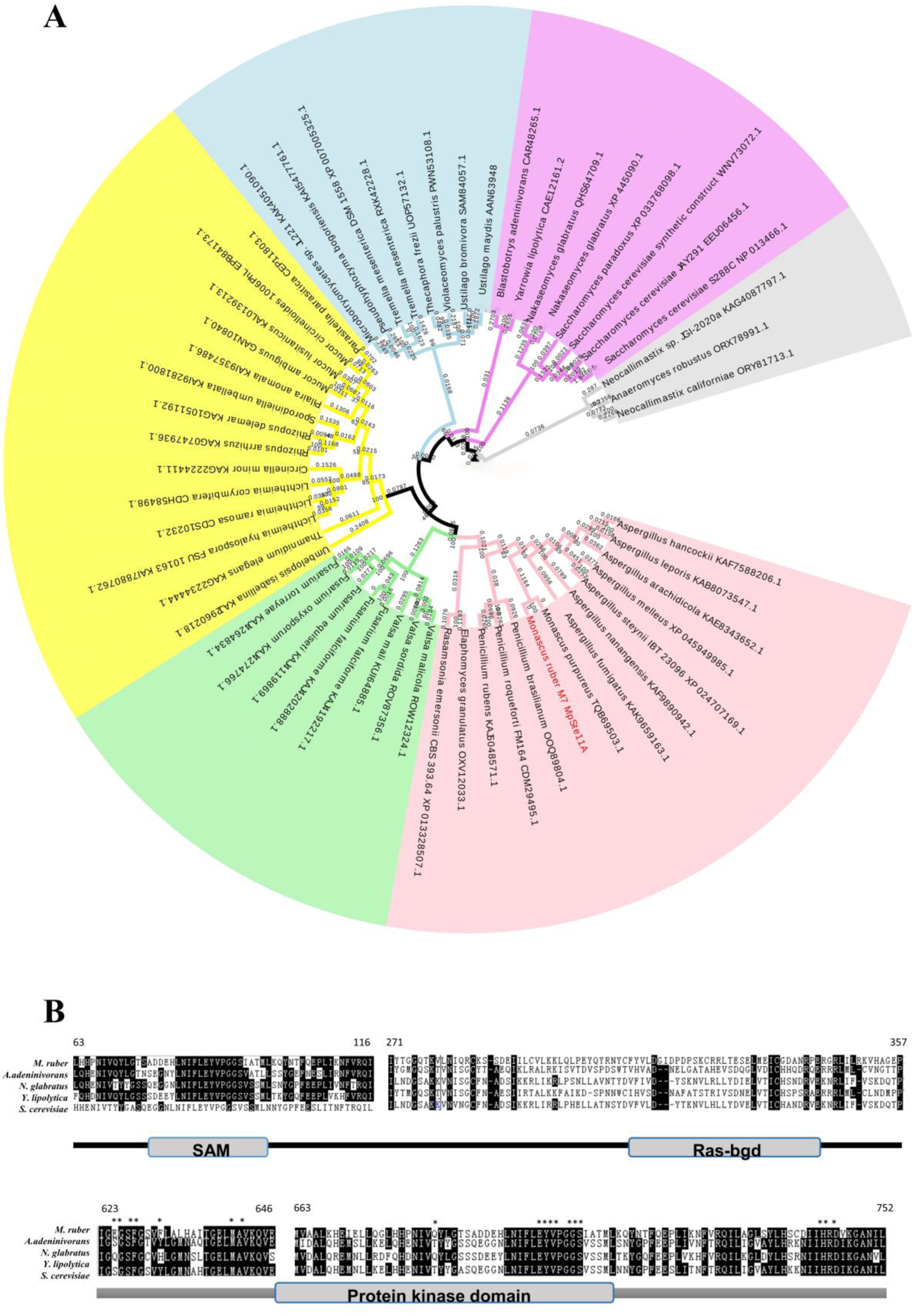
Bioinformatic analysis of fungal STE11 relative protein. (A) Phylogenetic tree of amino acid sequences of MpSte11A and various Ste11-like proteins, with color coding based on taxonomic classification: blue for *Basidiomycotina*, purple for *Saccharomycotina*, grey for *Mastigomycotina*, pink for *Ascomycotina*, and yellow for *Zygomycotina*. (B) Sequence comparison of Ste11-like protein kinase domains, *ATP binding site.

We aligned the protein sequence of MpSte11A with those of another four STE11 homologs derived from distinct fungal species (Fig. 1B). The results showed that MpSte11A contained characteristic SAM, Ras-binding, and protein kinase domains, exhibiting high homology with known STE11 proteins(26). As a member of the serine/threonine protein kinase family, STE11 typically featured a conserved kinase domain(27). The SAM domain mediated protein-protein interactions essential for regulating various biological processes(28), and the Ras-binding domain interacted with Ras or Ras-like proteins through intermolecular *β*-sheet formation(29). The protein kinase domain is critical for signaling and regulating development, differentiation, and intercellular communication(30). Based on the highest sequence coverage (93%), complete domain architecture (SAM-RBD-Kinase), and phylogenetic clustering with characterized STE11 orthologs, MpSte11A was prediected to be the primary candidate STE11 for functional characterization in this study.

### Knockout and Overexpression of *mpSte11A*

To characterize the function of *mpSte11A*, we constructed *mpSte11A* knockout and overexpression vectors by a seamless cloning approach. The Δ*mpSte11A* and OE::*mpSte11A* mutant strains were subsequently obtained through *A. rhizogenes*-mediated transformation of *Monascus* spp. Sequencing results confirmed the correctness of the mutant strains, which would be utilized in further experiments.

The deletion of mpSte11A resulted in a significant reduction in the growth rate of Monascus spp. The Δ*mpSte11A*, OE::*mpSte11A*, and wild-type *M. ruber* M7 (WT) strains were inoculated onto PDA medium and cultured at 28 ℃ (Fig. 2A). On day 7, Δ*mpSte11A* displayed significantly deeper coloration compared to the WT, suggesting that MpSte11A might control secondary metabolite production in *M.ruber* M7. After 3 days, the average colony diameter of Δ*mpSte11A* was 0.6 cm, whereas the WT measured 1 cm (Fig. 2B). Furthermore, hyphal of WT colonies was dense, while that of Δ*mpSte11A* colonies exhibited sparse and loose hyphae. These findings indicated that the *mpSte11A* gene played an important role in the trophic growth of *Monascus* spp.

**FIG 2.**
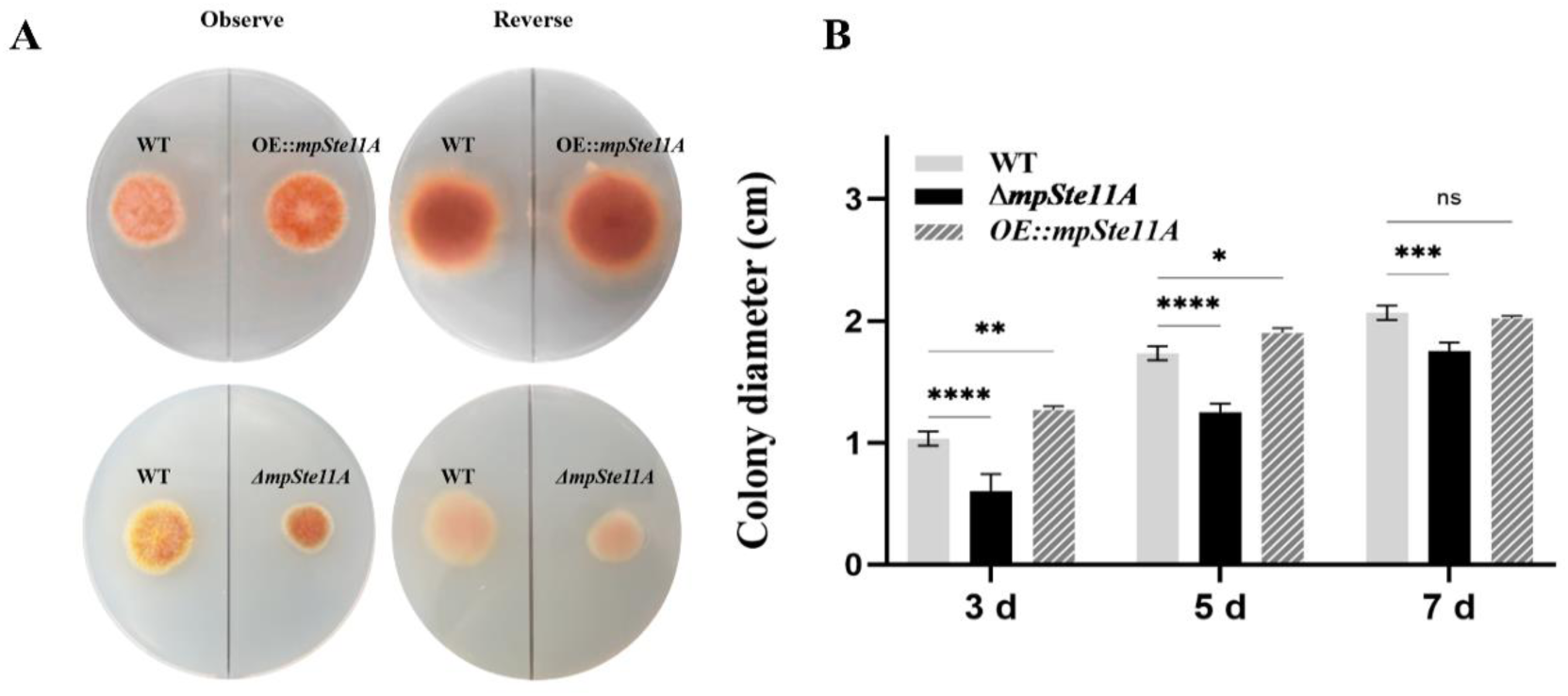
MpSte11A promoted the growth of *Monascus* spp. (A) The colony morphology of Δ*mpSte11A*, OE::*mpSte11A* and WT was observed after they were grown on PDA medium for 5 days. (B) Changes in colony diameter of Δ*mpSte11A*, OE::*mpSte11A* and WT strains grown on PDA medium for 3, 5, and 7 days, respectively.

### Knockout of *mpSte11A* Dramatically Enhancing MPs Production

To evaluate the role of MpSte11A in MPs biosynthesis, MPs production were compared among the Δ*mpSte11A*, OE::*mpSte11A*, and WT strains (Fig. 3). The WT strain reached a maximum biomass of 4.18 g/L on day 9, whereas Δ*mpSte11A* reached its peak on day 6 at 4.9 g/L. In contrast, OE::*mpSte11A* achieved the highest biomass of 6.0 g/L on day 9, which represented a 1.43-fold increase relative to the WT (Fig. 3A). These results indicated that deletion of MpSte11A promoted the mycelial growth of *M. ruber* M7.

**FIG 3.**
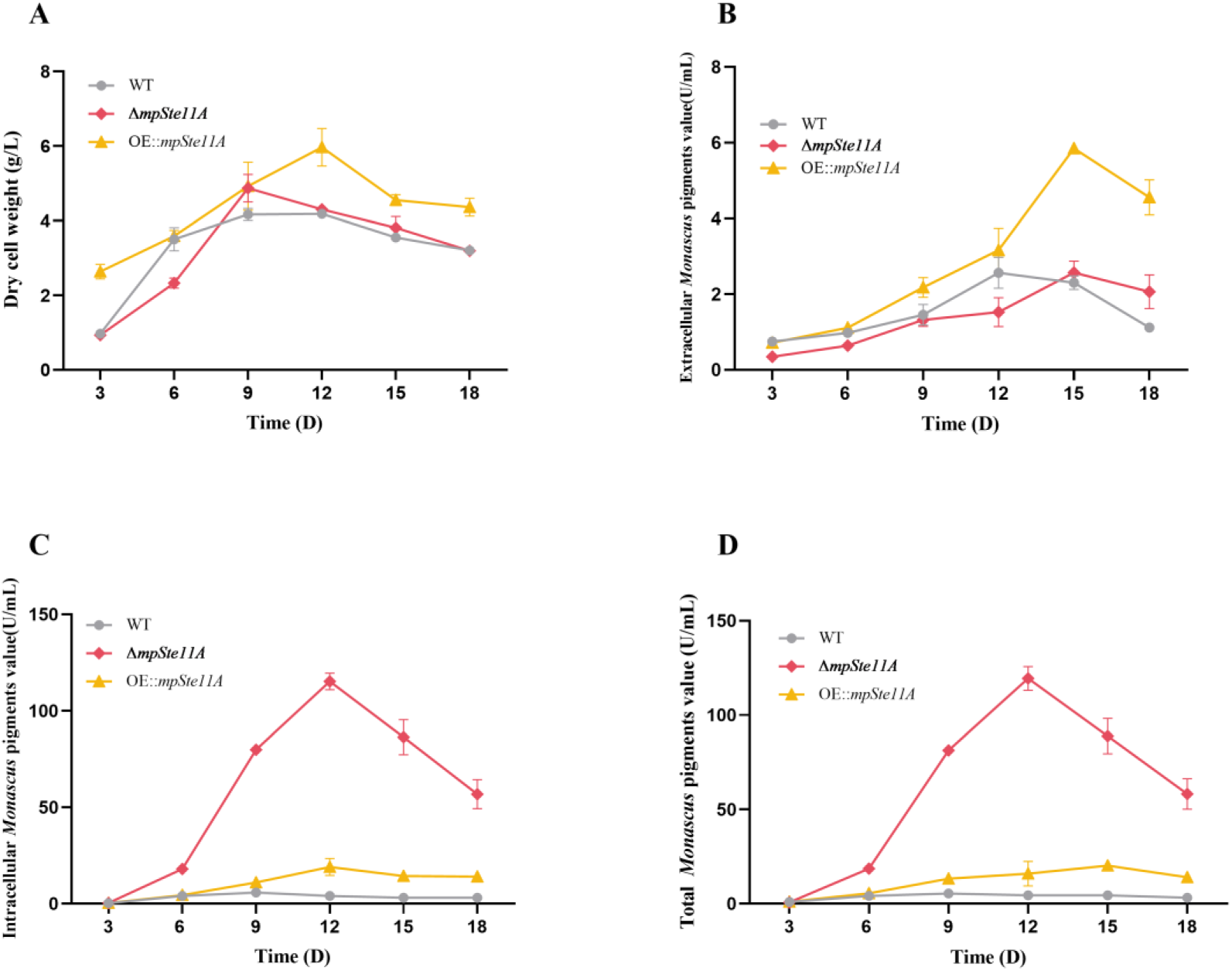
The Δ*mpSte11A* strain overproduced MPs. (A) Time-course changes in mycelial dry cell weight of the Δ*mpSte11A*, OE::*mpSte11A*, and WT strains. (B) Time-course profiles of total MPs accumulation in the Δ*mpSte11A,* OE::*mpSte11A*, and WT strains. (C) Time-course profiles of intracellular MPs accumulation in the Δ*mpSte11A,* OE::*mpSte11A*, and WT strains. (D) Time-course profiles of extracellular MPs accumulation in the Δ*mpSte11A*, OE::*mpSte11A*, and WT strains. All strains were cultured in PDB at 28 °C and 180 rpm for 18 days.

MPs are categorized into intracellular and extracellular fractions, and the total MPs is defined as the sum of both. Notably, the Δ*mpSte11A* strain reached a maximum intracellular MPs level of 119.4 U/mL on day 12, whereas the WT strain peaked at 5.5 U/mL on day 9. The intracellular MPs level of the Δ*mpSte11A* strain was approximately 22-fold higher than that of the WT (Fig. 3B). There were no significant differences in the production of extracellular MPs among the strains (Fig. 3C). Collectively, these results indicated that MpSte11A functioned as a negative regulator of MPs biosynthesis, and its deletion leads to an approximately 22-fold increase in total MPs production (Figure 3D).

### MpSte11A Controlling Hyphal Branching and Mycelial Growth

To investigate the role of MpSte11A in mycelial growth in *Monascus* spp., we compared the macroscopic size of the pellets and the hyphal morphology among the different mutant and the WT strains. We collected mycelial pellets of Δ*mpSte11A*, OE::*mpSte11A*, and the WT strains on day 5 for pellet diameter determination and scanning electron microscopy (SEM) analysis. The WT strain formed relatively large mycelial pellets with a rounded and smooth surface and a moderately compact texture. The Δ*mpSte11A* strain produced bright red mycelial pellets with a rough surface and exhibited a loose and soft texture. In contrast, the OE::*mpSte11A* strain formed smaller and smooth surface with a comparatively firm texture (Fig. 4A). Quantitative analysis showed that the OE::*mpSte11A* strain produced significantly smaller mycelial pellets than those of the WT and Δ*mpSte11A* strains (Fig. 4B).

**FIG 4.**
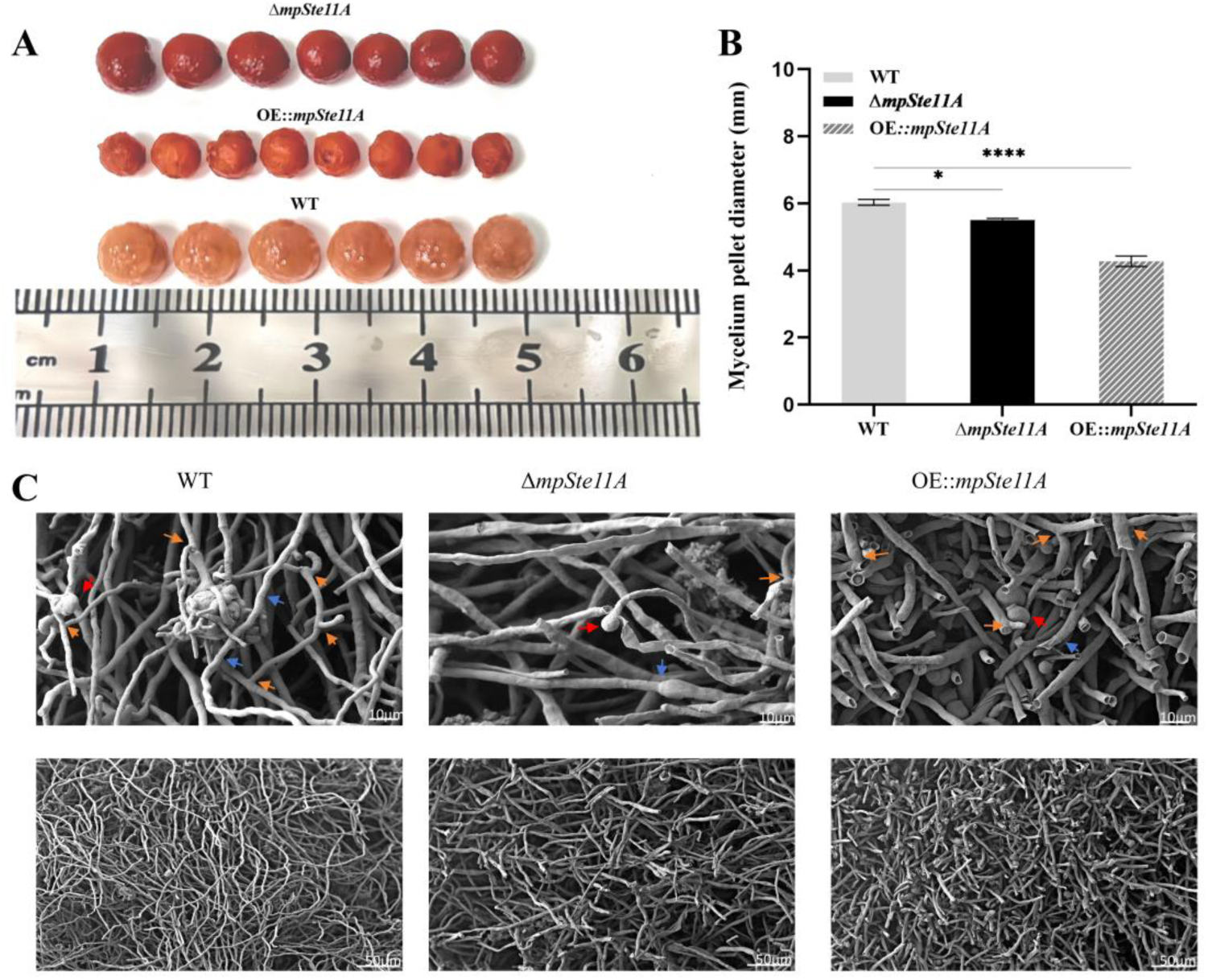
MpSte11A significantly controlled mycelial growth. (A) Comparison of mycelial pellet size between recombinant strains and the WT. (B) Statistical analysis of mycelial pellet diameters for recombinant strains and the WT. (C) SEM image of the hyphase in mycelial pellet. Mycelial pellets were fixed in 10 mL of 2.5% (v/v) glutaraldehyde at 4°C for 12 h. The internal morphology of pellets was examined by SEM at magnifications of 300× and 1000×.

In terms of microscopic morphology, the WT strain showed the rounded and smooth hyphae which formed a highly entangled and densely aggregated network. In contrast, the hyphae of Δ*mpSte11A* strain appeared dry, flattened, and irregular morphology. In the OE::*mpSte11A* strain, the hyphae were relatively rounded and plump with clearly defined hyphal branching points. These observations demonstrated that MpSte11A played a significant role in regulating hyphal branching and development in *Monascus* spp.

We also found that MpSte11A played a crucial role in regulating reproduction in *Monascus* spp. Deletion of *mpSte11A* completely abolished cleistothecia formation but markedly increased conidial production. In contrast, overexpression of *mpSte11A* slightly promoted cleistothecia formation but strongly suppressed conidiation. These results indicated that MpSte11A positively promoted sexual reproduction but inhibited asexual reproduction (Fig. S1). It was found that hyphal branching was reduced by 40% in Δ*mpSte11A* compared to the WT strain, with an average of only 4 hyphal branches per 500 μm (Fig. 5). In addition, we found differences in hyphal development between Δ*mpSte11A*, OE::*mpSte11A*, and WT strains. Our study demonstrated that *mpSte11A* regulated the growth and development of *Monascus* spp. and was essential for its hyphal branching.

**FIG 5.**
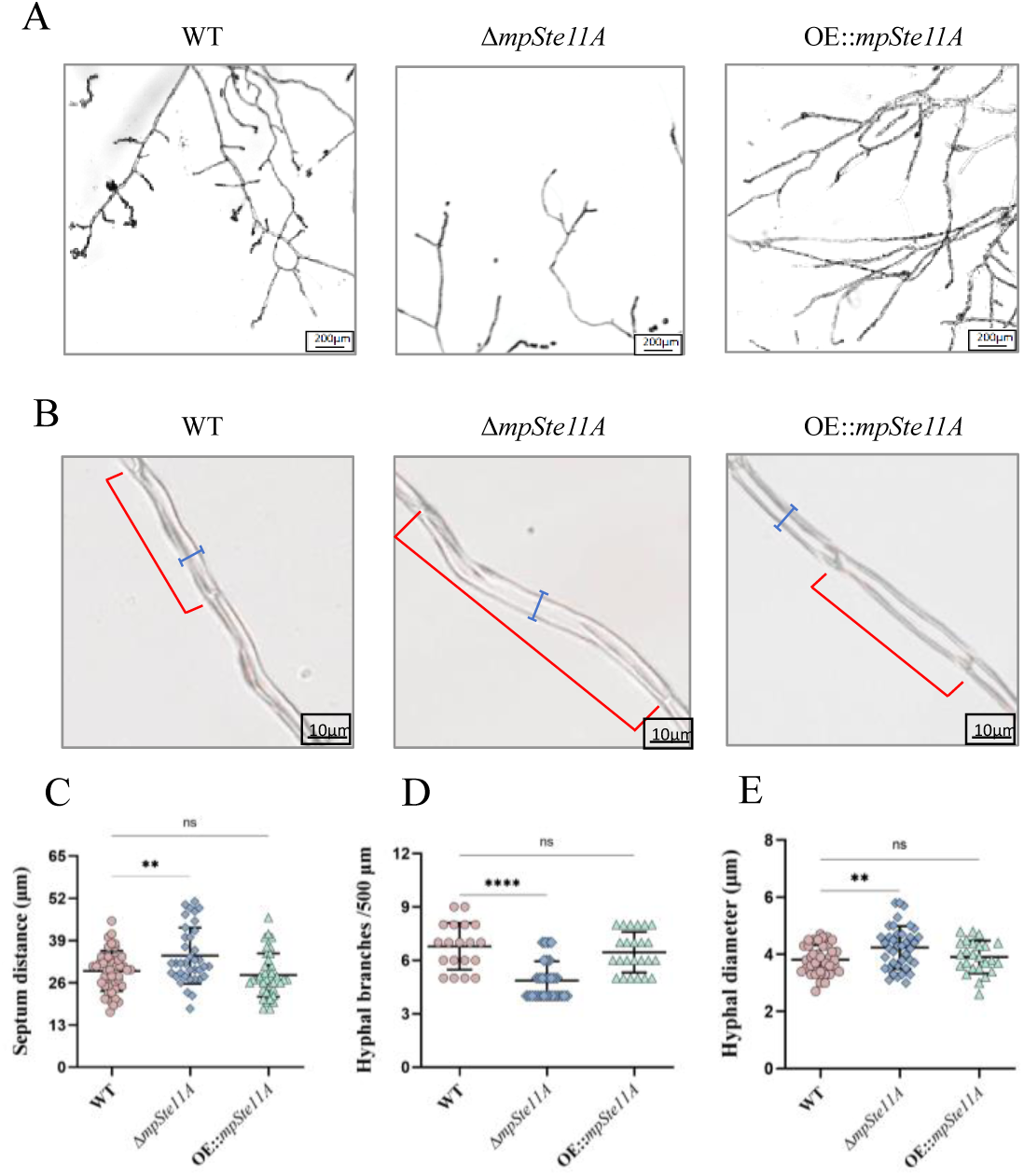
MpSte11A regulated hyphal branching and development. (A) Hyphal branching differences in recombinant strains and the WT strains. (B) Hyphal septum distance and hyphal diameter in recombinant strains and the WT. (C) Septum distances for the WT and mutant strains. (D) Number of hyphal branches per 500 μm for the three strains. (E) Hyphal diameters for the three strains. Culture conditions: Cultured statically on PDA medium at 28°C for 9 days, repeat the experiment 3 times.

### Joint transcriptomics and metabolomics analyses revealing MpSte11A redirecting Carbon Flux toward MPs biosynthesis

To explore the reasons for the phenotypes observed in strain Δ*mpSte11A*, we performed integrated transcriptomic and metabolomic analysis of the mutant and WT strains. This multi-omics approach identified 703 differentially expressed genes (DEGs) and 56 differentially regulated metabolites (DRMs). Among them, 480 genes were up-regulated and 223 gene were down-regulated in the Δ*mpSte11A* strain (Fig. S2-S3). Strikingly, most DEGs were up-regulated, the majority of DRMs (43 out of 56) were reduced, which suggested that increased transcriptional activity did not result in metabolites accumulation. KEGG pathway enrichment analysis highlighted five major perturbed pathways: starch and sucrose metabolism, TCA cycle, amino acid metabolism, ABC transporters, and polyketide biosynthesis (Fig. 6). Among the DRMs, 11 key metabolites met stringent criteria (Fragmentation Score >60, VIP >1.0, *P* <0.05) for significant depletion, including critical TCA cycle intermediates (succinate, isocitrate, α-ketoglutarate) and fatty acids (palmitic acid), which indicated a profound disruption of primary carbon metabolism.

**FIG 6.**
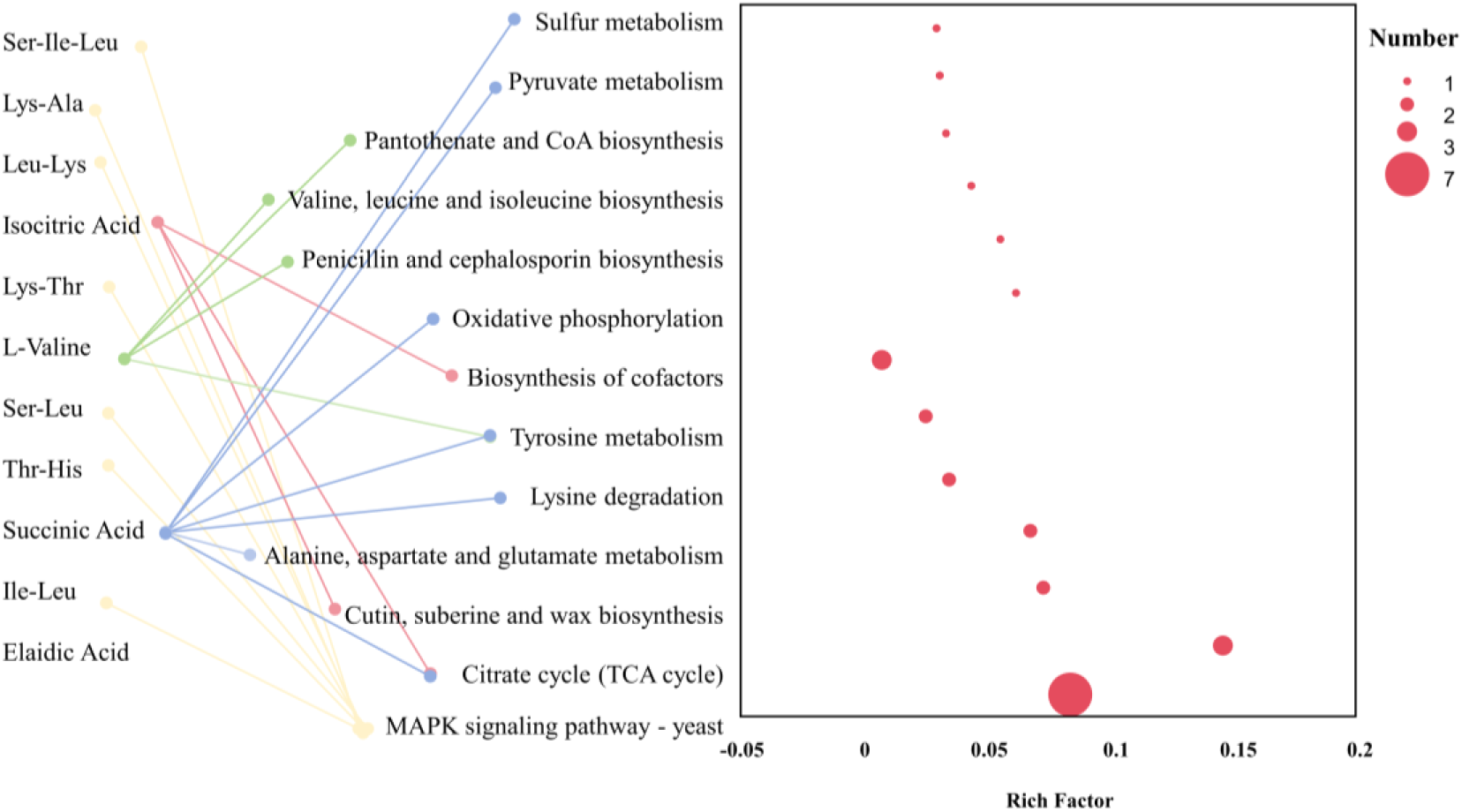
KEGG Enrichment Analysis of DEGs Corresponding to DRMs. KEGG pathway enrichment analysis of DEMs. The x-axis shows the Rich factor (Sample number/Background number), indicating the enrichment degree. The dot size represents the number of DEMs in the pathway.

Integration of transcriptomic and metabolomic data revealed that MpSte11A functioned as a metabolic gatekeeper whose deletion triggered a cascade of metabolic rearrangements (Fig. 7). The 22-fold increase in MPs production in Δ*mpSte11A* reflected a fundamental redirection of carbon flux from primary to secondary metabolism. When the TCA cycle was severely restricted, acetyl-CoA, a universal biosynthetic precursor, accumulated and was diverted into alternative pathways. A pyruvate decarboxylase (*PDC*) was down-regulated, which might shift pyruvate metabolism away from ethanol production toward acetyl-CoA supply for MPs biosynthesis.

**FIG 7.**
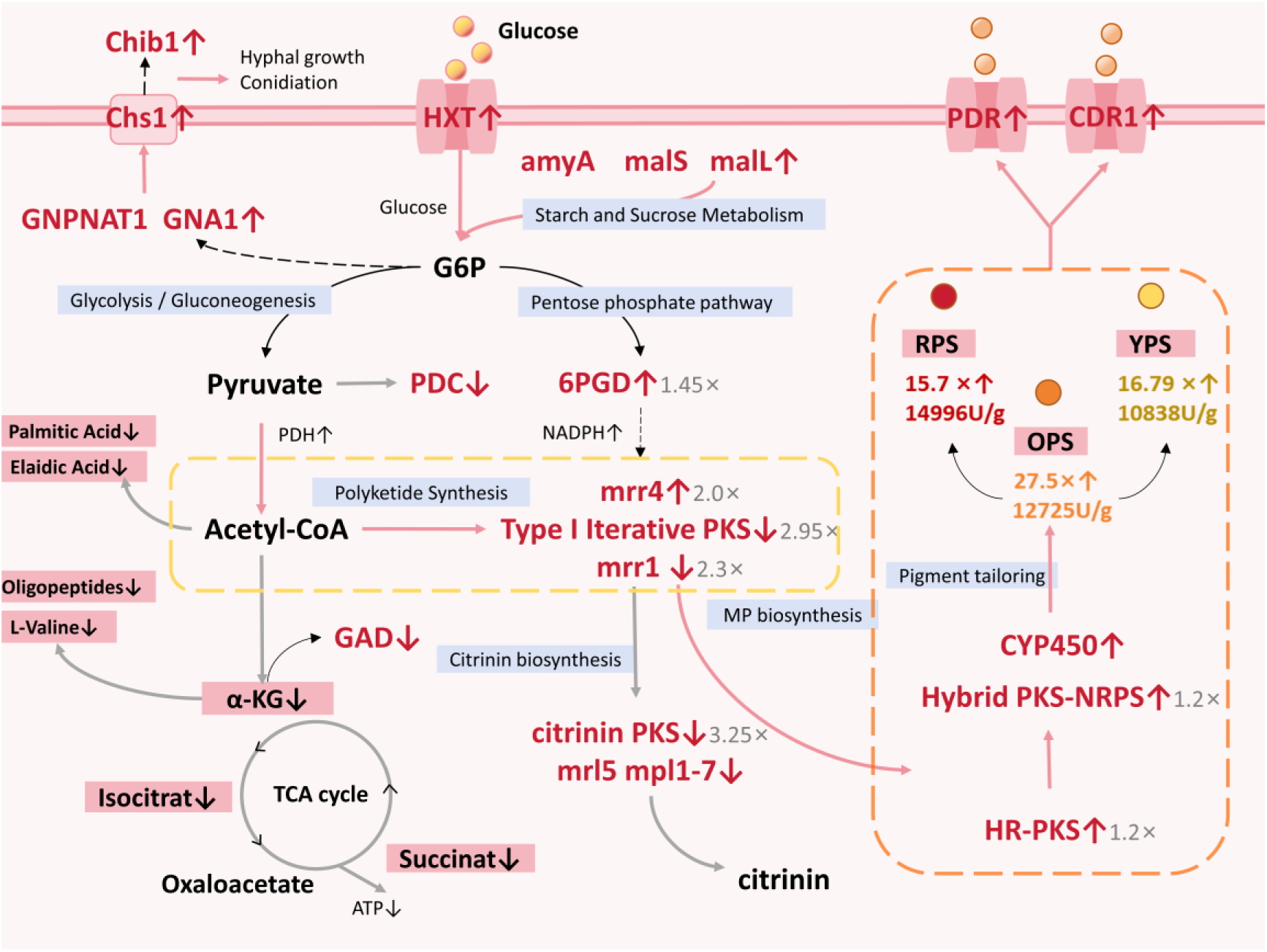
Deletion of *mpSte11* led to Metabolic Reprogramming. Metabolic pathways and differentially regulated metabolites are highlighted in blue and red boxes, respectively. Compounds are shown in black font, meanwhile differentially expressed genes and their encoded enzymes are labeled in red. For regulatory process, dashed lines indicate indirect regulation, thick gray lines represent inhibition, and thick pink lines denote promotion. The arrow direction corresponds to the up- or down-regulation of DRMs and DEGs.

Integrated multi-omics analysis established MpSte11A as a central metabolic coordinator whose activity determined the balance between primary and secondary metabolism in *Monascus* spp. From an applied perspective, MpSte11A acted as a key regulator of MPs production, and its deletion redirected cellular resources toward MPs biosynthesis, which offered a practical strategy for metabolic engineering of industrial *Monascus* strains.

## Discussion

In this study, we identified and characterized MpSte11A, a conserved MAPKKK in *Monascus* spp., and revealed its key role as a negative regulator of MPs biosynthesis. While MAPK signaling pathways are traditionally studied in the context of stress response, cell wall integrity, and pathogenicity in filamentous fungi(31–33), our integrated multi-omics analysis highlights a novel function of MpSte11A as a metabolic gatekeeper. The major finding was that deletion of *mpSte11A* triggered metabolic reprogramming. This redirected carbon flux from primary growth pathways toward overproduction of MPs, resulting in a 22-fold increase in intracellular MPs yield.

The dramatic enhancement of MPs production in the Δ*mpSte11A* strain could be explained by a shift in precursor availability. Our metabolomic data revealed a significant reduction of key TCA cycle intermediates, specifically succinate, isocitrate, and *α*-ketoglutarate, in the Δ*mpSte11A* strain. This suppression of the TCA cycle suggested a bottleneck in central carbon metabolism, which likely led to an accumulation of the upstream acetyl-CoA pool(34). Acetyl-CoA served as the central node linking primary energy metabolism and secondary polyketide biosynthesis(35). In the WT strain, MpSte11A appeared to direct acetyl-CoA primarily into the TCA cycle to promote hyphal growth and biomass accumulation. However, this primary metabolic sink was restricted in the Δ*mpSte11A* strain. Simultaneously, the downregulation of a pyruvate decarboxylase observed in our transcriptomic data suggested a reduction in the ethanol fermentation pathway, further driving pyruvate toward acetyl-CoA generation(36). This created a powerful force supplying excess cellular acetyl-CoA and malonyl-CoA toward the polyketide synthase (PKS) pathway, thereby promoting massive pigment biosynthesis(37). This phenomenon aligned with the overflow metabolism strategy, in which central carbon flux was redirected to secondary metabolites when downstream primary pathways were saturated or inhibited(38, 39). Traditional strain improvement often focused on overexpression of biosynthetic genes, but this strategy can be limited by precursor supply. Our results showed that manipulating regulators like MpSte11A could remove metabolic bottlenecks and redirect cellular resources more effectively.

The metabolic reprogramming in Δ*mpSte11A* was intimately coupled with alterations in fungal morphology. This mutant exhibited a transition from large and dense pellets to smaller and looser mycelial aggregates with reduced hyphal branching. While hyper-branching was often associated with high secretion capacity in some filamentous fungi(40), the loose pellet morphology observed here might offer a distinct advantage for intracellular MPs production. In submerged fermentation, dense pellets often suffered from hypoxic cores due to oxygen mass transfer limitations(41). The loose mycelial pellets of the Δ*mpSte11A* strain likely enhanced oxygen and nutrient diffusion to internal hyphae(42). Since MPs biosynthesis was an oxygen-consuming process(43), this morphological change supported the high metabolic demand of the mutant cell. Thus, MpSte11A likely regulated cytoskeletal organization and metabolic gene expression(44), and its removal shifted the cell into a state favorable for MPs production.

In summary, MpSte11A was not merely a developmental regulator but a central metabolic switch. Its deletion facilitated a targeted redirection of carbon flux away from the TCA cycle and into the MPs pathway. This mechanism of metabolic reprogramming provided a theoretical foundation for engineering overproducing fungal strains via modulation of MAPK signaling networks.

## Materials and Methods

### Microbial Strains, Plasmids and Chemical Regents

The strain used in this study was *M. ruber* M7 (CCAM 070120, Culture Collection of State Key Laboratory of Agricultural Microbiology, Wuhan, China)(45). Recombinant plasmids were constructed using *Escherichia coli* DH5α, and then *Agrobacterium tumefaciens* EHA105 was used as the plasmid DNA donor for the genetic transformation of *M.ruber* M7. Restriction endonucleases were obtained from Thermo Fisher Scientific (Waltham, USA)(6).

### Culture Medium and Growth Conditions

The mutant strains *ΔmpSte11A*, OE::*mpSte11A* and wild-type strain *M. ruber* M7 were inoculated in Czapek yeast extract broth (CYB) medium at 28℃ for 7 days and then the spores were washed with sterile water to obtain a suspended conidial solution, and the quantity was adjusted to 10^5^/mL. 400μLof spore solution was aspirated in 100 ml of Potato dextrose broth (PDB) medium at 28℃ and 150 rpm for 7 days.

### Construction of Knockout Strain Δ*mpSte11A* and Overexpression Strain OE::*mpSte11A*

The knockout strain *ΔmpSte11A* and overexpression strain OE::*mpSte11A* were constructed by homologous recombination, and the *mpSte11A* gene (2871 bp) as well as the 5’flanking region (1147 bp) and 3’flanking region (1100 bp) were amplified by using primers designed from the *Monascus ruber* M7 genome, the *gpdA* gene (1024 bp) was used as a promoter, and *neo* (1221 bp) was amplified from the pKNI plasmid as a marker for screening *ΔmpSte11A*. Cloned DNA fragments were ligated to plasmid pCAMBIA3300 (digested with *Xbal*I and *Hind* III) using pEASY®-Basic Seamless Cloning and Assembly Kit (TransGen Biotech, Beijing, China) to form plasmids.Recombinant plasmids were obtained by transformation screening of *Escherichia coli* DH5α and *A. tumefaciens* EHA105, and finally mutant strains of *Monascus ruber* M7 were screened by *A. tumefaciens* mediated transformation method.

### Morphology of Strains

The spore solution adjusted to 10^5^/mL was spot-applied 2.5 μL to PDA plates and incubated at 28℃ for 7 days, and the differences in colony morphology between *M.ruber* M7 and *ΔmpSte11A* and OE::*mpSte11A* were observed photographed and the colony diameters were measured on days 3, 5, and 7, respectively. Some sterile slide were inserted after 100 μL of spore solution was applied in the PDA plate and incubated at 28℃ for 3 days to observe the microscopic morphological changes.

### Determination of *Monascus* Pigments

The color values of the mycelium and the bacterial liquid were determined according to the established method with appropriate modifications. After extracting the MPs by weighing 20 mg of mycelium powder and diluting it with 80% (v/v) ethanol for appropriate times, the yellow MPs, orange MPs and red MPs were detected at 380 nm, 470 nm and 520 nm by UV-visible spectrophotometer, and the color valence of the mycelium and the pigmentation of the bacterial liquid were calculated, respectively.

### Transcriptomics Analysis

RNA purification, reverse transcription, library construction and sequencing were performed at Shanghai Majorbio Bio-pharm Biotechnology Co., Ltd. (Shanghai, China) according to the manufacturer’s instructions. Constructs of knockout strain *ΔmpSte11A*, overexpression strain OE::*mpSte11A*, and original strain *M.ruber* M7 were cultured in PDB medium at 28℃ for 6 days at 150 rpm, mycelia were collected, chilled with liquid nitrogen for 30 minutes, and stored at -80℃ for transcriptomics analysis. The analysis was performed by Majorbio Bio-pharm Technology (Shanghai,China).

### Metabolite Extraction

Fifty milligrams of solid sample were added to a 2 mL centrifuge tube along with a 6 mm diameter grinding bead. To extract metabolites, 400 μL of extraction solution (methanol:water = 4:1, v:v) containing 0.02 mg/mL of the internal standard (L-2-chlorophenylalanine) was added. The samples were ground for 6 minutes at -10°C and 50 Hz using a Wonbio-96c frozen tissue grinder (Shanghai Wanbo Biotechnology Co., Ltd.), followed by low-temperature ultrasonic extraction for 30 minutes at 5°C and 40 kHz. After incubation at -20℃ for 30 minutes, the samples were centrifuged for 15 minutes at 4°C and 13,000 g. The supernatant was then transferred to an injection vial for LC-MS/MS analysis.

### Quality Control Sample

As a part of the system conditioning and quality control process, a pooled quality control sample (QC) was prepared by mixing equal volumes of all samples. The QC samples were disposed and tested in the same manner as the analytic samples. It helped to represent the whole sample set, which would be injected at regular intervals (every 5-15 samples) in order to monitor the stability of the analysis.

## CRediT authorship contribution statement

**Tingya Wang**: Data curation, Validation, Writing - original draft; Investigation, Methodology, Writing - original draft; Formal analysis, Writing - original draft, Writing - review & editing; **Yali Duan**: Investigation; **Yuting Liu**: Writing; **Mu Li**: Project administration, Supervision, Funding acquisition, Writing - review & editing.

## Abbreviations

MPs: *Monascus* pigments
CYB: czapek yeast extract broth
PDB: potato dextrose broth
DRMs: differentially regulated metabolites
DEGs: differentially expressed genes
MAPK: mitogen-activated protein kinase
KEGG: Kyoto encyclopedia of genes and genomes
SEM: scanning electron microscopy

## Acknowledgments

This work was financially supported by the National Natural Science Foundation of China (Grant No. 32572534) and the Fundamental Research Funds for the Central Universities (Grant No. 2662024SZ003).

